# Hunchback functions in the post-mitotic larval MDN to restrict axon outgrowth, synapse formation, and backward locomotion

**DOI:** 10.1101/2025.07.31.667971

**Authors:** Kristen Lee, Natalie Rico Carvajal, Josmarie Graciani, Chris Q Doe

## Abstract

During neurodevelopment, a single progenitor cell can generate many different neuron types. As these neurons mature, they form unique morphologies, integrate into neural circuits, and contribute to behavior. However, the integration of these developmental events is understudied. Here, we show that the same transcription factor is important for both the generation of neuronal diversity and maintaining mature neuronal identity, providing novel insights on how the generation of neuronal identity and morphology are coordinated. We utilized a previously characterized larval locomotor circuit in *Drosophila*, where activation of the Moonwalker Descending Neuron (MDN) triggers backward locomotion via its presynaptic connection with the premotor neuron A18b. MDN expresses the temporal transcription factor Hunchback (Hb), which has a well-characterized role in neural progenitors. Loss of Hb in the post-mitotic MDN increases axon/dendrite branching, leading to additional functional synapses on A18b and increasing backward locomotion. We conclude that the endogenous function of Hb is to restrain axon/dendrite outgrowth, including limiting MDN-A18b synapses, thereby dampening backward locomotion. Our work provides insights on how a transcription factor can have different functions throughout life – i.e. Hb generates neuronal diversity in the progenitor and regulates neuronal connectivity in the mature neuron to generate an appropriately tuned behavior.

## Introduction

A major goal of neuroscience is to understand the relationship between mechanisms that generate molecular neuronal diversity and those determining circuit assembly and behavior. Work over the past decade has shown that spatial patterning is used to specify distinct progenitor pools in mammals or even single progenitors (neuroblasts) in flies (Briscoe et al., 2000; Skeath and Thor, 2003). A second step in generating diversity within individual progenitor lineages is temporal pattering, in which successively born neurons acquire a distinct identity based on a transcription factor cascade (Doe, 2017; El-Danaf et al., 2023; Pollington et al., 2023). However, the coordination between the mechanisms generating neuronal diversity and the maintenance of neuronal circuity in a mature animal remains largely unexplored.

The best characterized temporal transcription factor (TTF) cascade is in the *Drosophila* embryonic ventral nerve cord (VNC; analogous to the mammalian spinal cord), where neuroblasts sequentially express Hunchback (Hb), Kruppel, Pdm1/2, and Castor (Cas) (Doe, 2017). Similarly, the Hb mammalian ortholog, Ikaros (Ikz1), and *C. elegans* ortholog, *hbl-1*, are involved in neural progenitor temporal patterning (Blackshaw and Cayouette, 2025; Lin et al., 2003). Each TTF is transiently expressed in progenitors/neuroblasts, and then inherited by neuronal progeny born during its expression window, in this way TTF expression can be maintained post-mitotically. While the role of Hb in fly and mouse progenitors has been well characterized (Alsio et al., 2013; Blackshaw and Cayouette, 2025; Hirono et al., 2012; Isshiki et al., 2001; Kanai et al., 2005; Novotny et al., 2002; Pearson and Doe, 2003; Tran et al., 2010), the role of Hb in larval or adult post-mitotic neurons is less well understood (Goto et al., 2011; Lee et al., 2022).

Although Hb expression in post-mitotic embryonic *Drosophila* VNC neurons does not alter motor neuronal identity (Hirono et al., 2012), Hb appears to function differentially in post-mitotic central brain descending interneurons. Recently, we have shown that Hb is expressed in the post-mitotic GABAergic Pair1 descending neuron, part of a larval backward locomotion circuit, and that Hb has a role in maintaining Pair1 synapse number, connectivity, and behavior (Lee et al., 2022). Interestingly, the Hb ortholog in *C. elegans*, hbl-1, is also required in the GABAergic DD motor neuron to regulate synapse number (Thompson-Peer et al., 2012). Additionally, in worms, flies, and mice, Hb (or its respective orthologue) is expressed in post-mitotic neurons with long projections (Alsio et al., 2013; Javed et al., 2023; Lee et al., 2022; Thompson-Peer et al., 2012). While these mammalian neurons are both excitatory and inhibitory, the post-mitotic function of Ikaros, the mouse Hb orthologue, remains unknown (Alsio et al., 2013; Javed et al., 2023). Taken together, these findings suggest that post-mitotic Hb regulation of synapses in long projecting neurons may be evolutionarily conserved. Here, we expand on this hypothesis by investigating the role of Hb in the post-mitotic Moonwalker Descending Neuron (MDN), a cholinergic excitatory long-projecting neuron in *Drosophila*, with the goal that our findings can be translated to mammals.

There are four MDNs in the brain, two in each brain lobe, which trigger backward locomotion when activated (Carreira-Rosario et al., 2018). Larval MDNs provide excitatory input to the Pair1 neurons, which induces a pause in forward locomotion when activated (Carreira-Rosario et al., 2018; Lee et al., 2022; Tastekin et al., 2018). In addition, the MDNs provide direct excitatory input to the A18b premotor neurons, which is a segmentally repeated premotor neuron in the VNC that is active during backward, but not forward, locomotion (Carreira-Rosario et al., 2018). While this is a simplified rendition of the larval backward locomotor circuit, extensive work has been done to characterize the neuronal circuits contributing to backward crawling – including intersegmental feedback circuits in the VNC (Kohsaka et al., 2019) and incorporation of circuits responding to averse stimuli (Omamiuda-Ishikawa et al., 2020). Although these studies are extremely valuable, they lack an interrogation of the molecular mechanisms mediating the development and maintenance of these circuits.

Here we focus on the role of Hb within the larval MDNs. We show that Hb is required to limit MDN neurite outgrowth and prevent MDN from forming functional ectopic synapses with its circuit partners, thereby revealing that Hb functions to prevent abnormally robust backward locomotor behavior. Understanding if Hb is functioning similarly among post-mitotic central brain neuronal cell types will provide important information about how early developmental mechanisms can have long-term influences on neuronal identity.

## Results

### Hunchback is required in the larval MDN to restrain backward locomotion

Here we focus on a specific neuron in the backward locomotor circuit, MDN, which - like its partner neuron Pair1 - is born during the embryonic Hb temporal window (Figure 1A) and maintains Hb expression throughout larval life (Figure 1B). There are two pairs of MDNs in the larval brain, and all four MDNs express Hb. To determine whether Hb may also be functioning in post-mitotic MDNs, we knocked down Hb specifically in larval post-mitotic MDNs and assayed locomotor behavior. Fortunately, the larval MDN split-Gal4 driver turns on around 4 hours after larval hatching, ensuring all our manipulations are in the post-mitotic MDN. We previously showed that our *UAS-Hb^RNAi^* transgene was highly efficient at knocking down Hb to low levels both pan-neuronally and in the Pair1 neuron (Lee et al., 2022), and we find the same strong Hb knockdown when expressing *UAS-Hb^RNAi^* in the MDN neuron (Figure 1C).

**Figure 1.**
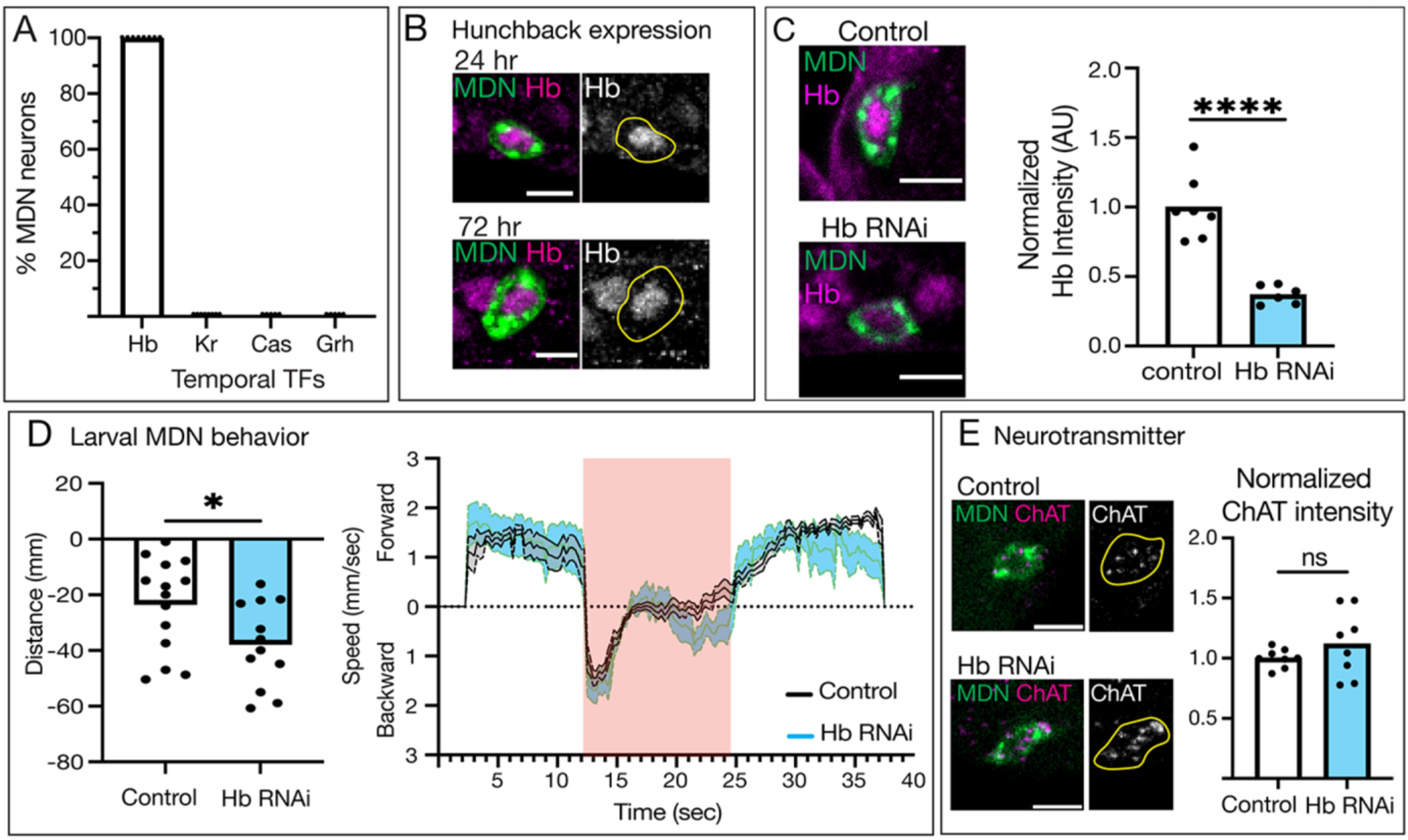
Hunchback is required in the larval MDN to restrain backward locomotion. **A)** Percent of MDNs expressing Hunchback (Hb), Krüppel (Kr), Castor (Cas), and Grainy head (Grh) in 0-2 hour old larvae, n = 8-12 animals. **B)** Hunchback (magenta, composite images; white, greyscale images) expression in MDN (green, composite images; yellow outline, greyscale images) at 24 and 72 hours after larval hatching. Genotype: *UAS-myr::smGdp-HA; GMR22H02_AD; GMR23E07_DBD.* Scale bar = 5µm. **C)** Compared to controls (white bar), Hb RNAi (blue bar) in MDN (composite image, green) significantly reduces but does not eliminate Hb expression (composite image, magenta). Control genotype: *UAS-myr::smGdp-HA; GMR22H02_AD; GMR23E07_DBD, UAS-Luc RNAi TRiP.JF01355*. Hb RNAi genotype: *UAS-myr::smGdp-HA; GMR22H02_AD; GMR23E07_DBD, UAS-Hb RNAi TRiP.HMS01183.* Scale bar = 5µm. Statistics: t-test, p < 0.0001, n = 6-7 animals. **D)** Left panel. Distance traveled (positive = forward distance, negative = backward distance) during optogenetic activation of control (white) and Hb RNAi (blue) animals. Statistics: t-test, p = 0.0336, n = 12-14 animals. Right panel. Directional speed over time before, during, and after optogenetic activation via a red-light stimulus (red square) of control (grey) and Hb RNAi (blue) animals. Control genotype: *UAS-CsChrimson::mVenus; GMR22H02_AD; GMR23E07_DBD, UAS-Luc RNAi TRiP.JF01355*. Hb RNAi genotype: *UAS-CsChrimson::mVenus; GMR22H02_AD; GMR23E07_DBD, UAS-TRiP.HMS01183 Hb RNAi.* **E)** Choline acetyltransferase (ChAT) expression (magenta, composite images; white, greyscale images) expression in MDN (green, composite images; yellow outline, greyscale images) at 72 hours after larval hatching does not change between control (white bar) and Hb RNAi (blue bar) animals. Control genotype: *UAS-myr::smGdp-HA; GMR22H02_AD; GMR23E07_DBD, UAS-Luc RNAi TRiP.JF01355*. Hb RNAi genotype: *UAS-myr::smGdp-HA; GMR22H02_AD; GMR23E07_DBD, UAS-Hb RNAi TRiP.HMS01183.* Statistics: t-test, p = 0.2677, n = 8 animals. Scale bar = 5µm.

In control animals, when the larval MDN is optogenetically activated, the animals move backward (Figure 1D). Interestingly, optogenetic activation of the larval MDN following Hb knock down promotes larvae to crawl backward a significantly longer distance than controls (Figure 1D, left panel), likely due to a second bout of backward locomotion during the optogenetic activation window (Figure 1D, right panel). We conclude that reducing Hb levels in the larval MDNs results in significantly more backward locomotion, suggesting that the normal function of Hb is to restrain the backward locomotor circuit.

We hypothesized that Hb is required to maintain aspects of larval MDN identity, similar to its function in the Pair1 neuron. First, we assayed neurotransmitter expression. MDN is an excitatory cholinergic neuron in the larval and adult (Bidaye et al., 2014; Carreira-Rosario et al., 2018), and we use Hybridization Chain Reaction (HCR) to assay RNA levels of choline acetyltransferase (ChAT) in control and Hb knockdown animals (Figure 1E). We opted to use in-situ hybridization for these experiments because the ChAT antibody was unreliable. When Hb was knocked down, larval MDN still expressed ChAT at similar levels as controls (Figure 1E), showing that Hb is not regulating MDN neurotransmitter identity. We conclude that Hb restrains MDN-induced backward locomotion, but is not required for MDN neurotransmitter choice.

### Hunchback restrains larval MDN axon/dendrite branching

Hb is not required for axon or dendrite morphology in the larval Pair1 neuron (Lee et al., 2022). To see if Hb regulates axon or dendrite morphology in larval MDN neurons, we knocked down Hb and labeled single MDNs using Multi-Color Flip-Out (MCFO) (Nern et al., 2015) (Figure 2). We used the image analysis software Imaris to trace each neuron (Figure 2A “Filaments”), allowing us to quantify the branching and length of dendrites and axons. The MDN has an ipsilateral dendrite and crosses the midline to form a contralateral dendrite before sending a descending axon to segment A4 in the VNC (Figure 2A; Carreira-Rosario et al., 2018). When Hb was knocked down, both the ipsilateral and contralateral dendrites had significantly more branches (Figure 2A-D) and an increased summed neurite length (Figure 2F, G). Similarly, there were significantly more axon branches and total length (Figure 2A, B, yellow arrow; 2E, H).

**Figure 2.**
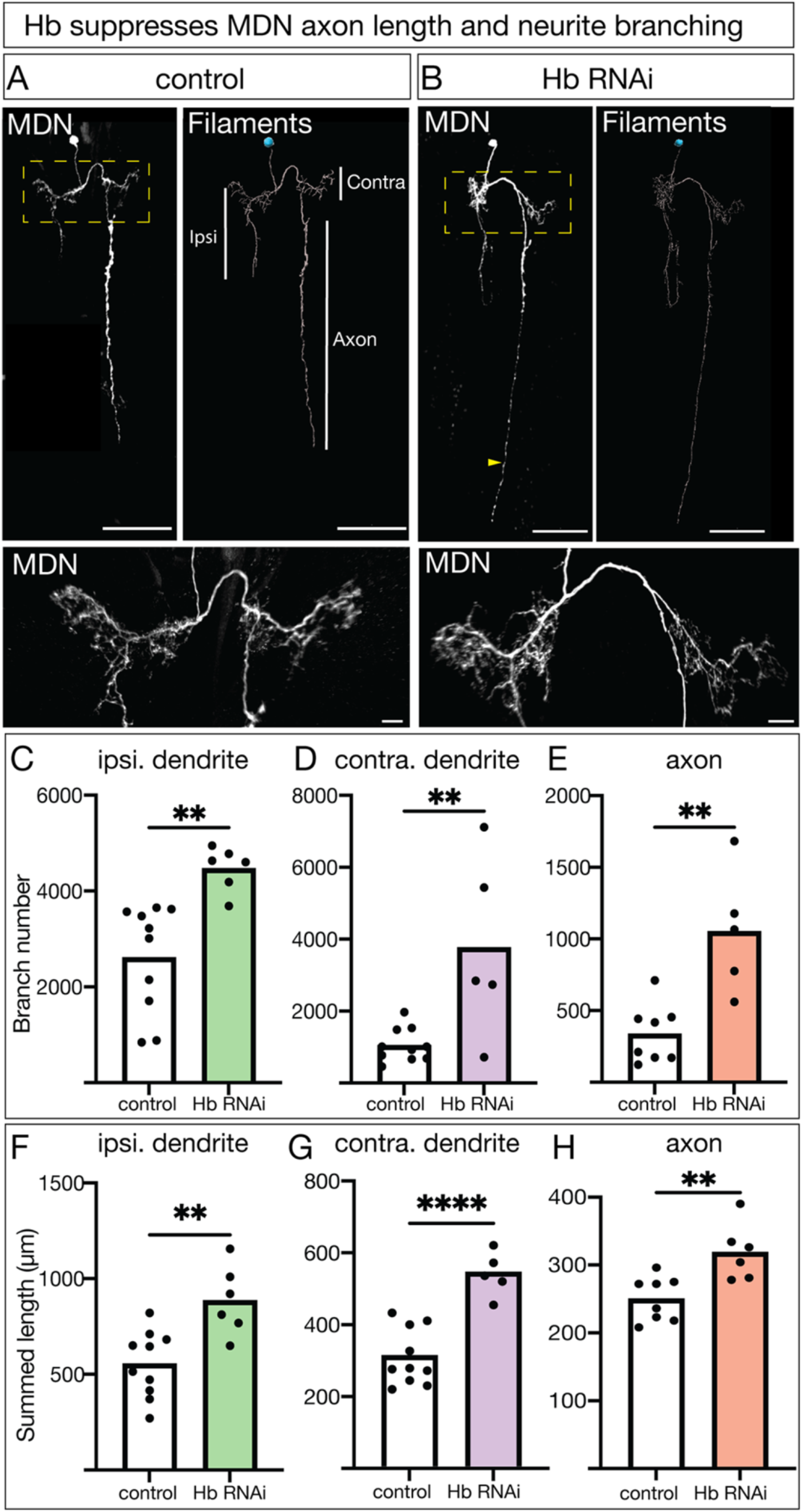
Hunchback restrains larval MDN axon/dendrite branching. **A, B)** Representative images of single labeled MDN in control (**A**) and Hb RNAi (**B**) animals. *In vivo* (left) and Imaris image analysis software tracing (filaments, right) shown. Scale bar = 50µm. Yellow dashed box around zoomed in dendritic region, scale bar = 10µm. Yellow arrow highlighting extended axon projection. The ipsilateral (ipsi.) dendrite is on the same side of the cell body, the contralateral (contra.) dendrite is on the opposite side from the cell body, and the axon descends into the ventral nerve cord on the contralateral side. **C-E)** Branch number for the ipsilateral dendrite (**C**), contralateral dendrite (**D**) and axon (**E**) in control (white) and Hb RNAi animals (colored bars). Statistics: (**C**) t-test, p = 0.0019, n = 6-10 neurons; (**D**) t-test, p = 0.0045, n = 5-10 neurons; (**E**) t-test, p = 0.0017, n = 5-8 neurons. **F-H)** Summed length of all traced neurites of the ipsilateral dendrite (**F**), contralateral dendrite (**G**), and axon (**H**) in control (white) and Hb RNAi animals (colored bars). Statistics: (**F**) t-test, p = 0.0027, n = 6-10 neurons; (**G**) t-test, p < 0.0001, n = 5-10 neurons; (**H**) t-test, p = 0.0045, n = 6-8 neurons. Control genotype: hs-Flp-G5::Pest; 10X UAS(frt.Stop)myr::smGDP-V5-THS-10XUAS(frt.Stop)myr::GDP-Flag/ *GMR22H02_AD*; *GMR23E07_DBD, UAS-Luc RNAi TRiP.JF01355.* Hb RNAi genotype: hs-Flp-G5::Pest; 10X UAS(frt.Stop)myr::smGDP-V5-THS-10XUAS(frt.Stop)myr::GDP-Flag/ *GMR22H02_AD*; *GMR23E07_DBD, UAS-Hb RNAi TRiP.HMS01183*.

We conclude that Hb is required in MDN to limit branch number and total axon/dendrite length. These results differ from our previous findings on Hb function in Pair1, but are consistent with a role for Hb in restraining backward locomotion described in the previous section.

### Hunchback prevents axon and pre-synapse targeting to lower abdominal segments

To characterize the effect of Hb knockdown on MDN morphology in more detail, we focused on its descending axon. The MDN-Gal4 driver labels two descending neurons: the MDN and a more medial Hb-negative neuron, which we have termed the Hb-negative descending neuron (DN; Figure 3A). Here we use the Hb-negative DN as an internal control - as it should not be affected by Hb knockdown, has a unique morphology from MDN, and both terminate in a similar location in the abdominal ganglion (Figure 3A, B). To confidently identify abdominal segments A1-A7, we used the segmentally repeated cluster of lateral Even-skipped (Eve) positive neurons (Figure 3C, D). In controls, the MDN descending axon invariably terminated in segment A4, although it occasionally extended slightly into segment A5, but never in segments A6-A7 (Figure 3C, white arrow; quantified in 3E); the same is true for the Hb-negative DN internal control neuron (Figure 3C, yellow arrow; quantified in 3E). In contrast, Hb knockdown resulted in the MDN axon fully extending into segments A5-A7 (Figure 3D, white arrow; quantified in 3E). Importantly, the Hb-negative DN control neuron still primarily targeted the A4 segment (Figure 3D, yellow arrow; quantified in 3E), thereby highlighting both the specificity of the Hb knockdown and its role in MDN axon mis-targeting.

**Figure 3.**
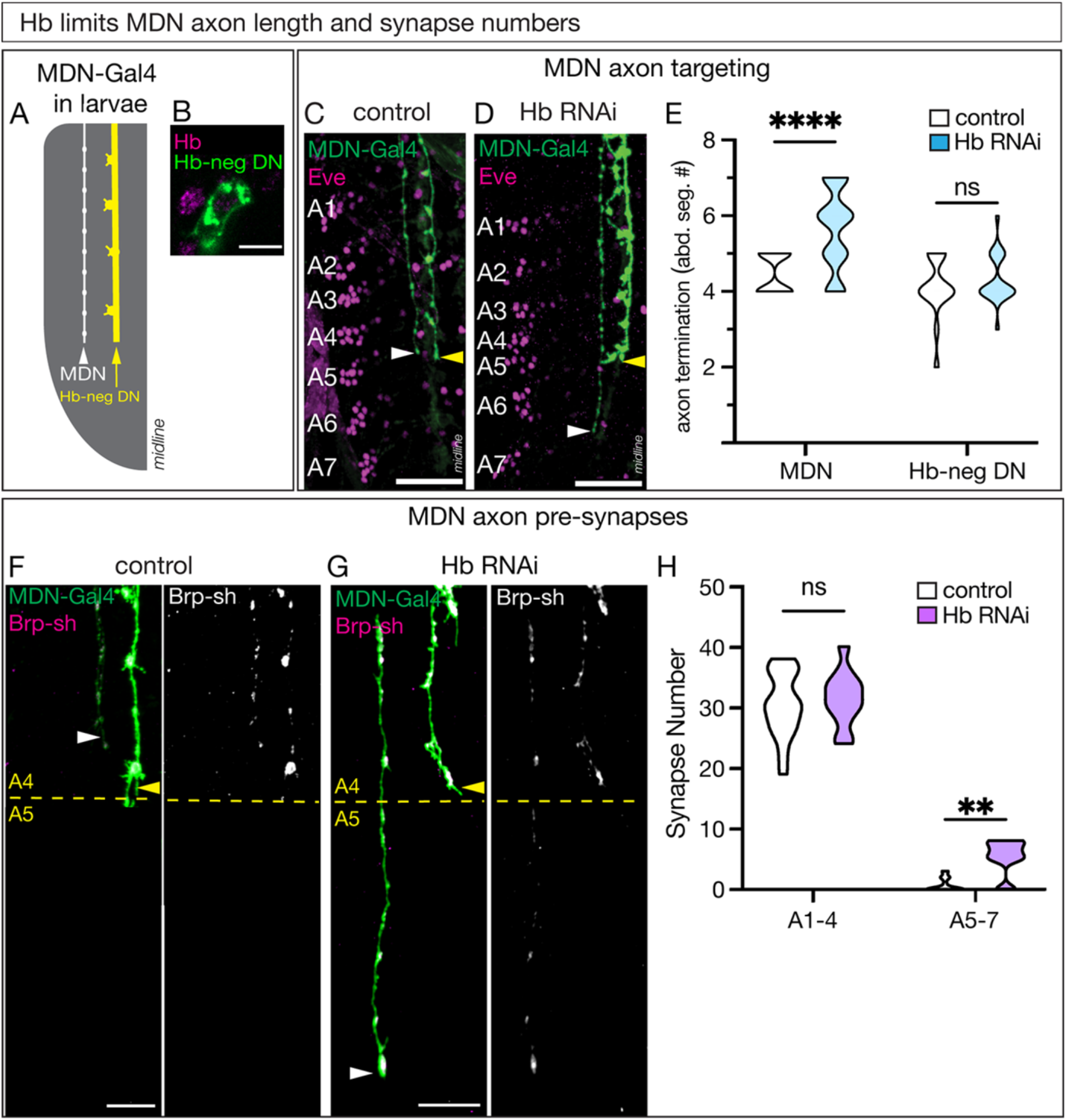
Hunchback is required for larval MDN axon targeting and synapse localization. **A)** Schematic of the SS01613-Gal4, hereafter referred to as “MDN-Gal4”, expression pattern in a hemi-segment of a third instar larval VNC. The MDN axon (white) is anatomically distinct from the Hunchback-negative off-target axon (yellow). **B)** The Hb-negative descending neuron (Hb-neg DN; green) does not express Hb (magenta) in the larvae. Scale bar = 5µm. **C-E)** MDN-Gal4 (green) showing the MDN axon (white arrow) and off-target axon (yellow arrow) in control (**C**) and Hb RNAi (**D**) animals. Even-skipped (eve; magenta) labeling abdominal segments A1-A7. Scale bar = 25µm. Quantification of abdominal segments the MDN (left) and off-target (right) axon terminates (**E**) in control (white) and Hb RNAi (blue) animals. Statistics: two-way ANOVA: genotype, F (1, 90) = 13, p = 0.0003; neuron, F (1, 90) = 20, p < 0.0001; interaction, F (1, 90) = 13, p = 0.0102; Bonferroni’s multiple comparisons between genotypes for each neuron: MDN, p < 0.0001; off-target, p = 0.9241; n = 18-38 VNC hemi-segments. Control genotype: *UAS-myr::smGdp-HA; GMR22H02_AD; GMR23E07_DBD, UAS-Luc RNAi TRiP.JF01355*. Hb RNAi genotype: *UAS-myr::smGdp-HA; GMR22H02_AD; GMR23E07_DBD, UAS-Hb RNAi TRiP.HMS01183.* **F-H)** MDN-Gal4 (green) showing the MDN axon (white arrow) and off-target axon (yellow arrow) in control (**F**) and Hb RNAi (**G**) animals. Labeling of pre-synapses in MDN-Gal4 pattern using Bruchpilot (Brp-sh; magenta, composite images). Yellow dashed line marks the boundary between the A4 and A5 segment designated by Eve staining. Scale bar = 10µm. (**H**) Quantification of the number of MDN pre-synapses anterior of A5 (A1-4, left) and posterior of A5 (A5-7, right) in control (white) and Hb RNAi (purple) animals. Statistics: two-way ANOVA: genotype, F (1,56) = 8.1, p = 0.0060; abdominal segment, F (1,56) = 757, p < 0.0001; interaction, F (1, 56) = 3.1, p = 0.0794; Bonferroni’s multiple comparisons between genotypes for each abdominal segment location: A1-5, p = 0.9032; A5-7, p = 0.0035; n = 12-18 VNC hemi-segments. Control genotype: *UAS-myr::smGdp-HA; GMR22H02_AD, UAS-brp-D3-mStrawberry; GMR23E07_DBD, UAS-Luc RNAi TRiP.JF01355*. Hb RNAi genotype: *UAS-myr::smGdp-HA; GMR22H02_AD, UAS-brp-D3-mStrawberry; GMR23E07_DBD, UAS-Hb RNAi TRiP.HMS01183*.

To determine if the MDN had pre-synapses in the ectopic A5-A7 axon extension, we used the MDN-Gal4 to express a non-functional presynaptic marker, Bruchpilot-short (Brp), and assayed the number of synapses on the MDN axon posterior or anterior to A5. This boundary was determined by Eve staining (Figure 3F, G, dashed line) and by the Hb-negative DN internal control axon which showed unchanged termination in A4 (Figure 3F, G, yellow arrow). In controls, the MDN axon primarily terminated at segment A4, as expected (Figure 3F), with Brp+ pre-synapses distributed along the axon (Figure 3F, Brp-sh). Hb knockdown showed no change in the number of pre-synapses in segments A1-A4; however, we detected Brp+ synapses in the ectopic posterior axonal domain (Figure 3G; quantified in 3H). We conclude that Hb acts in MDN to prevent ectopic synapse formation, either directly or indirectly through restraining axon extension. These findings are consistent with a role for Hb in restraining backward locomotion described above.

### Hunchback prevents MDN innervation of A18b dendritic regions in posterior abdominal segments

The larval MDN circuit is well-characterized (Carreira-Rosario et al., 2018; Lee and Doe, 2021; Lee et al., 2022) (Figure 4A). One neuron in this circuit is A18b, is a cholinergic pre-motor neuron downstream of MDN located in the VNC, which is active only during backward locomotion (Carreira-Rosario et al., 2018). Given that larval Hb knockdown leads to an *increase* in backward locomotion and MDN axon mistargets to abdominal segments A5-A7, we hypothesized that Hb knockdown may result in ectopic MDN-A18b synapses in A5-A7.

**Figure 4.**
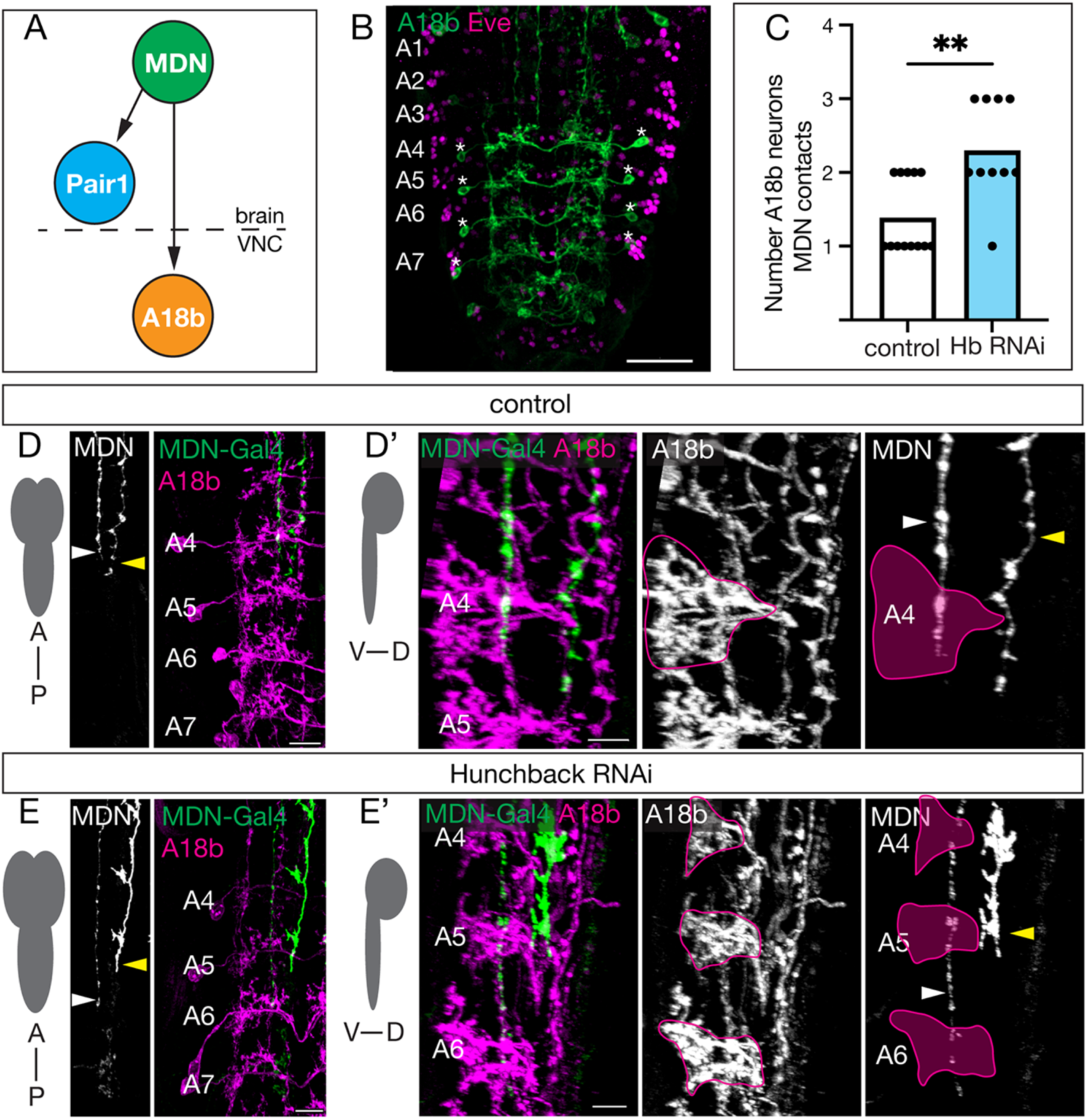
Hb prevents MDN innervation of A18b+ dendrite volume in A5-A7. **A)** Schematic of the core MDN larval circuit. MDN forms synapses with Pair1 in the central brain and A18b in the VNC. **B)** A18b neurons (green; asterisk marking cell body) are present in abdominal segments A4-A7 (Eve, magenta) at 72 hours after larval hatching. Scale bar = 30µm. **C)** Quantification of the number of A18b neurons the MDN axon contacts within the VNC at 72 hours after larval hatching in control (white bar) and Hb RNAi (blue bar) animals. Statistics: t-test, p = 0.0013, n = 10-13 VNC hemi-segments. **D/D’)** Region of MDN and A18b dendritic innervation in control animals. **D)** Anterior to posterior view of MDN-Gal4 (white, greyscale image; green, composite image) innervating A18b dendrites (magenta, composite image) in segments A4-7 in control animals at 72 hours after larval hatching. White arrow, MDN axon; yellow arrow, Hb-negative DN off-target axon. Scale bar = 10µm. **D’)** Ventral to dorsal view of MDN-Gal4 (green, composite image; white, greyscale image) innervating the A18b dendritic region (magenta, composite image; white, greyscale image; magenta outline for clarity) in the A4 segment of the VNC. White arrow, MDN axon; yellow arrow, Hb-negative DN axon. Scale bar = 10µm. Genotype: *UAS-myr::smGdp-HA, LexAop-myr::smGdp-V5; GMR22H02_AD, 94E10-LexA; GMR23E07_DBD, UAS-Luc RNAi TRiP.JF01355*. **E/E’)** Region of MDN and A18b dendritic innervation in Hb RNAi animals. **E)** Anterior to posterior view of MDN-Gal4 (white, greyscale image; green, composite image) innervating A18b dendrites (magenta, composite image) in segments A4-7 in Hb animals at 72 hours after larval hatching. White arrow, MDN axon; yellow arrow, Hb-negative DN axon. Scale bar = 10µm. **E’)** Ventral to dorsal view of MDN-Gal4 (green, composite image; white, greyscale image) innervating the A18b dendritic region (magenta, composite image; white, greyscale image; magenta outline for clarity) in the A4-A6 segments of the VNC. White arrow, MDN axon; yellow arrow, Hb-negative DN axon. Scale bar = 10µm. Genotype: *UAS-myr::smGdp-HA, LexAop-myr::smGdp-V5; GMR22H02_AD, 94E10-LexA; GMR23E07_DBD, UAS-Hb RNAi TRiP.HMS01183*.

To test this hypothesis, we first confirmed that A18b neurons are located in A4-A7 in the third instar larval VNC (Figure 4B, asterisk). Next, we asked whether Hb knockdown increased MDN innervation of the A18b dendritic volume in segments A4-A7 by quantifying the number of A18b neurons that the MDN axon physically contact (Figure 4C). In controls, MDN innervates the A18b dendritic volume primarily in segment A4, but occasionally also in segment A5 (Figure 4D, white arrow; quantified in 4C). In addition to segment A4, Hb RNAi resulted in a robust innervation of the A18b dendritic volume in more posterior segments A5-A7 (Figure 4E, white arrow; quantified in 4C). Importantly, the Hb-negative DN does not contact A18b (Figure 4D-E, yellow arrow) We conclude that Hb RNAi results in MDN extending its axon into the more posterior segments A5-A7, where the MDN axon innervates the A18b dendritic domains. This raised the possibility that Hb RNAi may lead to ectopic MDN-A18b synapse formation.

### Hunchback prevents ectopic MDN-A18b putative synapses in posterior abdominal segments

To test the hypothesis that MDN forms ectopic synapses with more posterior A18b neurons when Hb is knocked down, we first needed a reliable way to label MDN synapses. Due to genetic limitations, we could not label both MDN synapses and A18b membranes in control and Hb knockdown animals. Instead, we utilized a previously published method showing that axonal varicosities strongly correlates with synapse location (Sales et al., 2019). We validated this method by demonstrating that the non-functional presynaptic marker, Brp-short, colocalizes exclusively with MDN axonal varicosities (Figure 5A). To determine if we could unbiasedly label these varicosities, we increased the contrast of the axonal image and then used the “spots” function in Imaris image analysis software to label the varicosities. We found that this technique revealed a strong correlation between MDN axonal varicosities and the Brp-short signal, thus allowing us to confidently label MDN synapses (Figure 5A’).

**Figure 5.**
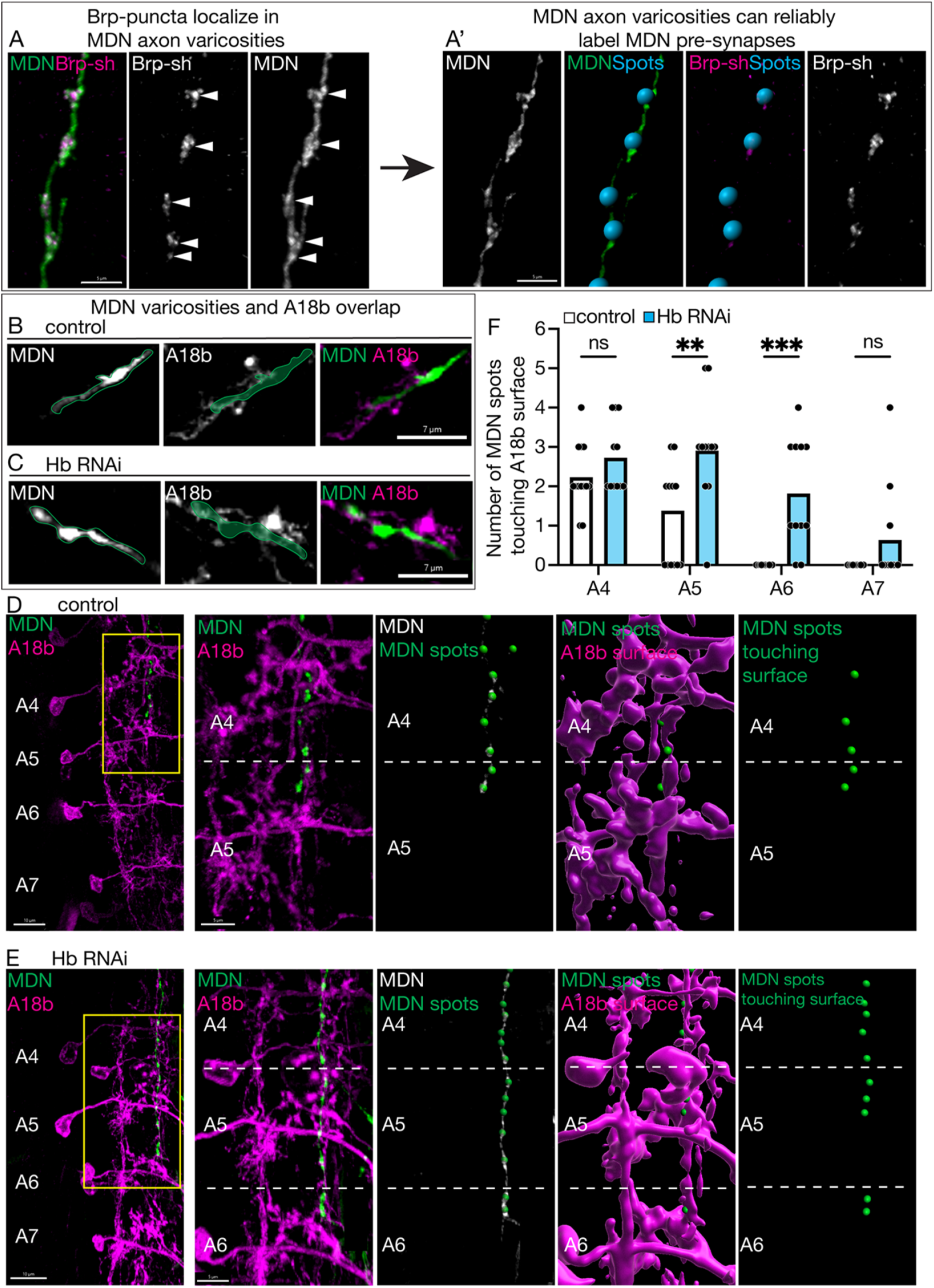
Hunchback prevents ectopic MDN-A18b putative synapses in posterior abdominal segments. **A/A’)** MDN axon varicosities can be used to label MDN pre-synapses. **A)** MDN axon (green, composite image; white, greyscale image) co-localized with Bruchpilot (Brp-sh; magenta, composite images; white, greyscale image; white arrows for location reference). Scale bar = 5µm. **A’)** MDN axon varicosities (white, greyscale image; green, composite image) were used as a reference for 3D Spots (blue, composite image). These Spots co-localized with Bruchpilot (Brp-sh; magenta, composite image; white, greyscale image). Scale bar = 5µm. **B, C)** Single slice images of MDN (white, greyscale image, green outline for reference; green, composite image) and A18b (white, greyscale image; magenta, composite image). **B)** control animals, **C)** Hb RNAi animals. Scale bar = 7µm. **D/D’)** Putative MDN synapses with A18b neurons in control animals at 72 hours after larval hatching. **D)** MDN putative synapses (green) and A18b neurons within abdominal segment A4-A7 (magenta). Scale bar = 10µm. Yellow box indicates zoomed in region in D’. **D’)** MDN putative synapses (green, composite image) and A18b neurons within abdominal segment A4 and A5 (magenta, composite image); dashed line separates A18b neuron in A4 from A5. Spots (green dot, composite image) indicates location of MDN putative synapse (white, composite image). MDN spots (green dot, composite image and standalone image) touching A18b surface (magenta, composite image) were assayed. Scale bar = 5µm. **E/E’)** Putative MDN synapses with A18b neurons in Hb RNAi animals at 72 hours after larval hatching. **E)** MDN putative synapses (green) and A18b neurons within abdominal segment A4-A7 (magenta). Scale bar = 10µm. Yellow box indicates zoomed in region in E’. **E’)** MDN putative synapses (green, composite image) and A18b neurons within abdominal segment A4-A6 (magenta, composite image); dashed line separates A18b neurons within unique segments. Spots (green dot, composite image) indicate location of MDN putative synapse (white, composite image). MDN spots (green dot, composite image and standalone image) touching A18b surface (magenta, composite image). Scale bar = 5µm. **F)** Quantification of the number of MDN putative synapse spots touching A18b neuronal surface in control (white bar) and Hb RNAi (blue bar) animals. Statistics: two-way ANOVA: genotype, F (1, 88) = 27, p < 0.0001; abdominal segment, F (3, 88) = 23, p < 0.0001; interaction, F (3, 88) = 2.3, p = 0.0777; Bonferroni’s multiple comparisons between genotypes within each abdominal segment: A4, p = 0.9830; A5, p = 0.0022; A6, p = 0.0002; A7, p = 0.5511; n = 11-13 VNC hemi-segments. Control genotype: *UAS-myr::smGdp-HA, LexAop-myr::smGdp-V5; GMR22H02_AD, 94E10-LexA; GMR23E07_DBD, UAS-Luc RNAi TRiP.JF01355*. Hb RNAi genotype: *UAS-myr::smGdp-HA, LexAop-myr::smGdp-V5; GMR22H02_AD, 94E10-LexA; GMR23E07_DBD, UAS-Hb RNAi TRiP.HMS01183*.

Next, we used this methodology to assay where MDN is forming synapses with A18b in control and Hb knockdown animals. First, we showed that MDN varicosities and A18b membranes overlap in a single optical slice (Figure 5B, C). Next, we used 3-dimensional data to determine how many synapses MDN was forming with each A18b neuron in abdominal segments A4-A7. We assayed MDN synapse number using axonal varicosities, as described above. To unbiasedly determine how many synapses MDN was forming with A18b, we used Imaris image analysis software to put a “surface” over the A18b dendritic domains. This allowed us to assay how many “spots” were touching a “surface”. In control animals, we found that MDN formed 3-6 total synapses with A18b neurons exclusively in the A4-A5 segments, with most residing in the A4 segment (Figure 5D; quantified in 5F). In contrast, Hb knockdown increased MDN synapses with A18b to an average of 7-12 total synapses between the A4-A7 abdominal segments (Figure 5E; quantified in 5F). There was a significant increase in MDN-A18b synapse number in segments A5 and A6 when Hb was knocked down (Figure 5F), which correlated with an increased likelihood that the MDN axon targeted these regions when Hb was knocked down (Figures 3, 4), demonstrating that MDN is forming ectopic synapses with A18b neurons. Therefore, Hb normally functions to limit the number of MDN-A18b synapses.

### Hunchback prevents ectopic MDN-A18b functional synapses in posterior abdominal segments

Next, we wanted to determine whether Hb knockdown resulted in *functional* synapses with A18b in A5-A7. To test this, we expressed a CaMPARI2 (Calcium Modulated Photoactivatable Ratiometric Integrator) transgene in A18b neurons (Moeyaert et al., 2018). CaMPARI functions as a photoconvertible protein, converting from green to red when exposed to 405nm illumination and concurrent elevation of calcium levels, i.e. neuronal activity (Figure 6A). The CaMPARI2 transgene specifically allows antibody detection of the original green (via anti-Flag) and photoconverted red (via anti-CaMPARI2) signals independently, allowing increased sensitivity when imaging. As previously described (Carreira-Rosario et al., 2018), we activated MDN via a noxious head poke, concurrently exposed the larvae to 405nm illumination, and assayed the number and segmental location of A18b neurons photoconverted to red in both control and Hb knockdown conditions (Figure 6B). As anticipated, all A18b neurons expressed the green CaMAPRI2 signal (Figure 6C, D, asterisk). In control animals, A18b neurons in segment A4, and occasionally segment A5, photoconverted to red, showing that MDN activation is causing increased activity in A18b neurons in this region (Figure 6C, white arrows; quantified in 6E). When Hb was knocked down in MDN, A18b neurons in the more posterior segments A5-A7 also photoconverted to red (Figure 6D, white arrows; quantified in 6E), showing that MDN activation is causing increased activity in A18b neurons in this region. A posterior neuron that is not A18b was also consistently activated in the Hb knockdown animals (Figure 6D, yellow arrow). Importantly, in both control and Hb knockdown animals, A18b was only photoconverted when a noxious head poke was administered, and not when the stimulus was absent (Figure 6E), showing that the experimental setup alone does not photoconvert A18b neurons to red. Taken together, these results show that MDN forms ectopic functional connections with A18b following Hb knockdown in MDN. Thus, when Hb is reduced, the MDN axon extends into more posterior segments but still prioritizes synaptic connections with neurons within its established locomotor circuit, its partner A18b (Figure 6F). We conclude that Hb is required in MDN to restrict the number and function of MDN-A18b synaptic contacts and thus dampen backward locomotor behavior.

**Figure 6.**
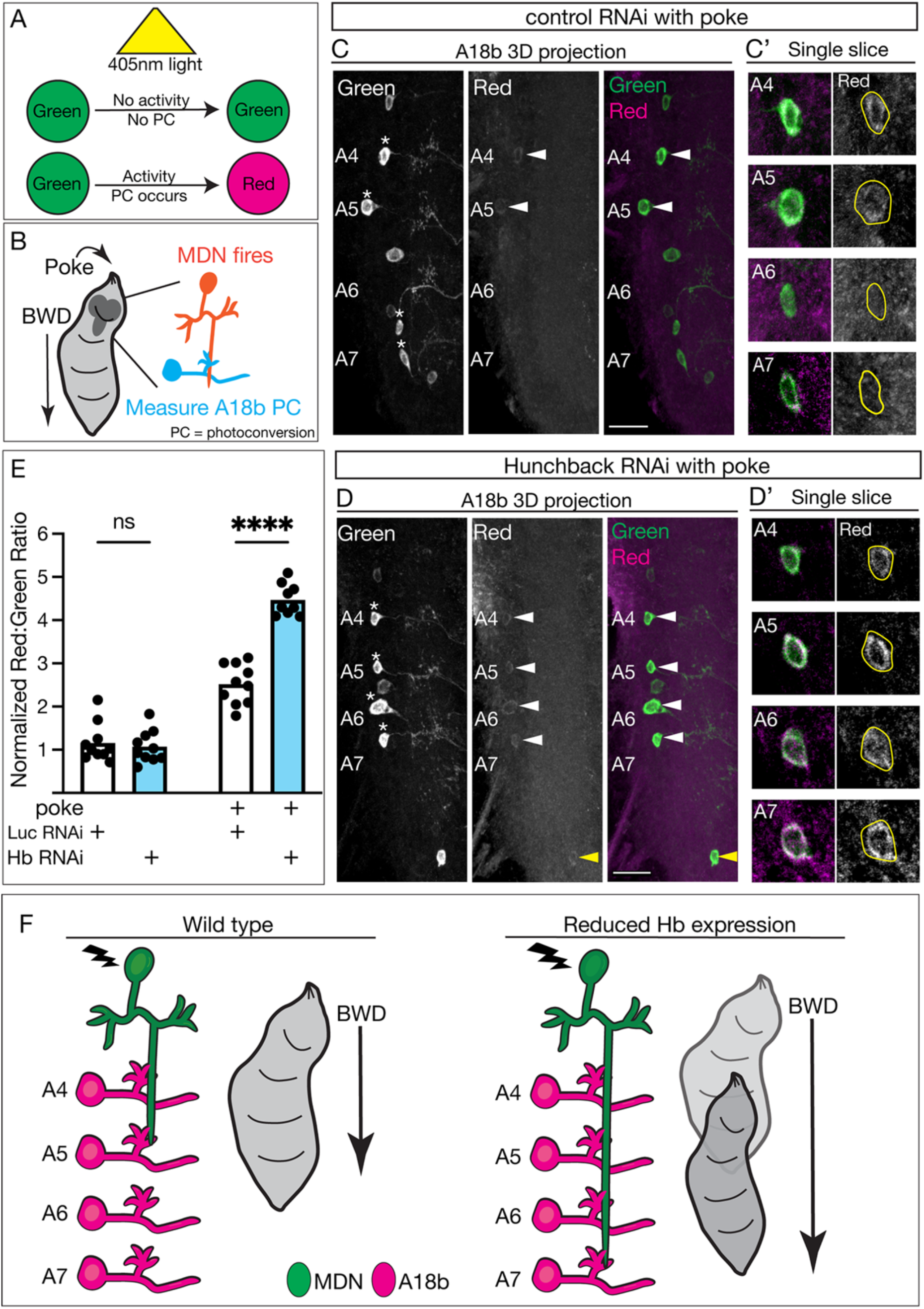
Hunchback prevents ectopic MDN-A18b functional synapses in posterior abdominal segments. **A)** Schematic of how the CaMPARI transgene determines whether a neuron experienced increased activity during exposure to photoconvertible light. If a neuron did not experience increased activity during the light exposure, it will remain green. If a neuron was active during the light exposure, it will photoconvert (PC) to red (depicted as magenta in figures henceforth). **B)** Schematic of the experimental protocol. A noxious head poke was used to manually induce backward (BWD) locomotion, causing MDN to fire. Photoconversion using the CaMPARI transgene was used to determine if the A18b neurons were activated in response to MDN activity. **C/C’)** A18b photoconversion during MDN activation (i.e. poke) in control animals at 72 hours after larval hatching. **C)** 3D projection of CaMPARI2 signal: all A18b cell bodies labeled green (marked with asterisk, white, greyscale image; green, composite image), and the cells that photoconverted to red (white, greyscale image; magenta, composite image). Abdominal segments A4-A7 labeled. White arrow shows positive CaMPARI2 PC signal. Scale bar: 15µm. **C’)** Single slices of A18 cell bodies in abdominal segments A4-A7 from the 3D projection image. Genotype: *LexAop-CaMPARI2; GMR22H02_AD, 94E10-LexA; GMR23E07_DBD, UAS-Luc RNAi TRiP.JF01355*. **D/D’)** A18b photoconversion during MDN activation (i.e. poke) in Hb RNAi animals at 72 hours after larval hatching. **D)** 3D projection of CaMPARI2 signal: all A18b cell bodies labeled green (marked with asterisk, white, greyscale image; green, composite image), and the cells that photoconverted to red (white, greyscale image; magenta, composite image) Abdominal segments A4-A7 labeled. White arrow shows positive CaMPARI2 PC signal in A18b. Yellow arrow shows positive CaMPARI2 PC signal in off-target neuron. Scale bar: 30µm. **D’)** Single slices of A18 cell bodies in abdominal segments A4-A7 from the 3D projection image. Genotype: *LexAop-CaMPARI2; GMR22H02_AD, 94E10-LexA; GMR23E07_DBD, UAS-Hb RNAi TRiP.HMS01183*. **E)** Quantification of the normalized red to green ratio of the lateral region of the VNC containing the A18b cell bodies at 72 hours after larval hatching in control RNAi (Luc RNAi; white bar) and Hb RNAi (blue bar) animals. Statistics: two-way ANOVA: genotype, F (1, 35) = 49, p < 0.0001; treatment, F (1, 35) = 320, p < 0.0001; interaction, F (1, 35) = 58, p < 0.0001; Bonferonni’s multiple comparisons: no poke control vs no poke Hb RNAi, p > 0.9999; poke control vs poke Hb RNAi, p < 0.0001; n = 9-10 VNC hemi-segments. **F)** Schematic of all results in wild type (right) and Hb knockdown (left) animals. MDN, green; A18b, magenta. Lightning bolt represents MDN activation.

## Discussion

Here we show that the MDN, a *Drosophila* central brain descending neuron required for initiating backward locomotion, is born in the Hb TTF window during neurogenesis. MDN expresses Hb throughout larval life, allowing us to assay its role in larval post-mitotic MDN neuronal identity maintenance. Larval MDN Hb knockdown results in increased backward distance traveled when MDN is optogenetically activated. We show that in control animals, MDN forms synapses with the pre-motor neuron A18b in abdominal segment A4. However, when Hb is knocked down, the MDN axon mistargets to the more posterior abdominal segments, A5-A7, and forms functional ectopic synapses with the A18b neurons in these segments. Thus, Hb restricts MDN-A18b synaptic contacts and dampens backward locomotor behavior.

These data add to a growing body of evidence suggesting that TTFs may be functioning differentially in the *Drosophila* larval central brain versus the VNC (Hirono et al., 2017; Lee et al., 2022). Although temporal patterning is better understood in VNC neuroblast lineages (Doe, 2017; Pollington et al., 2023), this work highlights the importance of studying the role of temporal patterning in the central brain, to better understand how this early developmental mechanisms can shape neuronal identity throughout life.

We were surprised to find that Hb is functioning differently in MDN and Pair1 neurons in the larval central brain. In Pair1 neurons, Hb does not alter morphology but is required for normal synapse number and behavior (Lee et al., 2022). While Hb is still required in MDN for synapse number, our major finding was that Hb is functioning to regulate axon outgrowth/guidance. This difference between Hb in MDN and Pair1 may be due to different Hb expression levels in these cells. Previous work has shown that Hb can be present in “high” or “low” quantities in a neuronal stem cell, leading to the generation of unique neuronal types (Gabilondo et al., 2011; Pearson and Doe, 2003). Additionally, larval MDN and Pair1 are located in different regions of the brain, enabling unique spatial factors to function in combination with temporal factors to maintain MDN morphology. Lastly, MDN and Pair1 express different neurotransmitters. Strikingly, both Pair1 in flies and the DD neuron in *C. elegans* are GABAergic, and do not have morphology changes when Hb/hbl-1 expression is reduced, instead showing synapse number changes (Lee et al., 2022; Thompson-Peer et al., 2012). Perhaps MDN’s excitatory profile, in conjunction with Hb expression, is contributing to its morphology changes. Indeed, the mammalian Hb ortholog, Ikaros, is expressed in both excitatory and inhibitory neurons in mice (Alsiö et al., 2013; Javed et al., 2023), suggesting that Ikaros may be differentially regulating morphology and synapse number post-mitotically in these cells as well.

During development, conserved axon guidance cues, i.e. attractive or repulsive signals, allow the axon to travel to the correct location within the nervous system (Evans and Bashaw, 2010). Mutation of the axon guidance cue Round-about (Robo) leads to abnormal behavior in larvae (Berni, 2015), demonstrating that axon pathfinding is important for whole animal function. The axon typically travels in multiple bouts of smaller distances between “guidepost” cells, before settling in a final location. Guidepost cells in insects, or intermediate targets in vertebrates, are specialized to provide local cues to either promote or inhibit the growth of a specific neuronal axon (Araújo and Tear, 2003). Although many genes have been implicated in axon pathfinding (Kraut et al., 2001), our work is the first to show that the TTF Hb is also involved in this process. We hypothesize that Hb may be required for the expression of an axon guidance cue, allowing the MDN axon to traverse the VNC and terminate on A18b in abdominal segment 4. Perhaps A18b neurons are acting as a guidepost cell for MDN axon pathfinding.

## Materials and Methods

### Fly husbandry

All flies were reared in a 25C room at 50% relative humidity with a 12 hr light/dark cycle unless noted otherwise. All comparisons between groups were based on studies with flies grown, handled and tested together.

### Fly stocks

- ;GMR22H02_AD (attp40); GMR23E07_DBD (attp2) (SS01613-Gal4) (Gift from J. Truman, University of Washington)
- LexAop-myr::smGdp-V5, UAS-myr::smGdp-HA;; (BDSC# 64092)
- ;; UAS-Hb RNAi TRiP.HMS01183 (attp2) (BDSC# 34704)
- ;; UAS-Luc RNAi TRiP.JF01355 (attp2) (BDSC# 31603)
- UAS-Chrimson::mVenus (attp8);; (BDSC# 55134)
- ; 10X UAS(frt.Stop)myr::smGDP-V5-THS-10XUAS(frt.Stop)myr::GDP-Flag; (BDSC# 62124)
- hs-FlpG5::Pest;; (BDSC# 77140)
- UAS-brp::sh-mStrawberry (BDSC# 80571)
- 94E10-LexA (attp40) (made in Doe lab)
- LexAop-CaMPARI2;; (Janelia #3028062)

### Immunohistochemistry

Standard confocal microscopy and immunocytochemistry methods were performed. In short, larval CNS were dissected in ice cold hemolymph-like buffer and fixed with 4% paraformaldehyde for 12-23 minutes, depending on age. The tissue was exposed to normalized donkey serum block for either 40 minutes at room temperature or overnight at 4C. After, the tissue was exposed to a primary antibody solution overnight at 4C. After being washed with 0.3% PBST, samples were exposed to a secondary antibody solution overnight at 4C. After a series of dehydration, tissue was mounted with DPX; the only exception was the CaMPARI experiments, which were mounted with ProLong Glass Antifade mountant (Thermo Fisher, Eugene, OR). Primary antibodies used: Rabbit anti-Hunchback (1:400; Doe lab), Rat anti-HA (1:100; Sigma #11867423001, St. Louis, MO), chicken anti-V5 (1:800; Bethyl A190-218A, Montgomery, TX), mouse anti-Flag (1:1000; Sigma F1804, St. Louis, MO), Rabbit anti-Eve (1:500; Doe lab), rabbit anti-DsRed (1:200; Abcam ab62341, Waltham, MA), mouse anti-CaMAPRI2 (1:1000; Absolute Antibody). Secondary antibodies were from Jackson ImmunoResearch (Donkey, 1:400; West Grove, PA).

### Image acquisition and processing

Confocal image stacks were acquired on a Zeiss 900 airyscan microscope using Nyquist sampling. All images were process either in Fiji (https://imagej.new/fiji) or Imaris (https://imaris.oxinst). Methodology for specific analyses using these software packages are described below. Figures were made using Adobe Illustrator.

### Quantification of Pixel Intensity

All pixel intensity quantification was done manually using the “Measure” feature in Fiji. The freehand tool was used to outline the cell body. Measurements were set to “Area” and “Raw Integrated Density”. A middle slice of the total cell body was measured. For each cell body, the “Raw Integrated Density” was divided by the “Area”. The sum of these values is reported.

### Behavior

Embryos were transferred to a food bottle and aged until 48 hr after larval hatching, then the larvae were transferred to apple caps with a food mixture of 0.5 µM All Trans Retinol (ATR) or 0.5 µM ethanol (sans ATR control), 6 mL of deionized water, and 5 mL of yeast grain for 24 hours. Behavior experiments were all done with third instar larvae aged 72-76 hr after larval hatching. Larvae were transferred to 1.3% agar sheets on the FIM table. FIM table uses frustrated total internal reflection (FTIR) to capture high-resolution and high-control videos for experiments. After a two-minute accumulation, larvae locomotion was recorded at 4 frames per second for a 150-frame video (37.5 seconds) with a Basler camera. At 50 frames, the red light was turned on for 50 frames (turned off at 100 frames). Next, the behavior video was uploaded to Fiji to crop the video and increase the contrast, creating a darker background. Then, it was uploaded to FIMTracker to track larval behavior. Directional information was acquired using the “mov_direction” calculations. Distance traveled and acceleration was reported.

### Multicolor FlpOut and quantification of morphology

Stage 16/17 embryos were heat shocked at 37C for 8-10 minutes in a water bath. After heat shock, embryos recovered at 18C for an equal amount of time. The CNS was dissected from 70-74hr old animals. Images of the brains were imported into the Imaris Image Analysis Software. The filament tool was used to trace the morphology of MCFO-labeled MDN. The largest diameter was set to 3.5 µm and the smallest diameter was 0.9 µm. Defining the primary neurite as the neurite extending from the cell body, the ipsilateral dendrite was the material ipsilateral to, but not including the primary neurite. The contralateral dendrite was the material contralateral to, but not including, the primary neurite. The primary neurite is the neurite extending from the cell body. The first projection off the descending neurite was used as a landmark for the beginning of the axon region and the end of the contralateral dendrite region. This landmark had been previously characterized (Carreira-Rosario et al., 2018).

### Abdominal segment identification

We used Even-skipped (Eve), which is expressed in a lateral cluster of 10 neurons per abdominal segment, to identify the A1-A7 segments. Boundaries were defined by the most posterior region of each Eve-positive cluster.

### Pre-synapse quantification with Brp-short

Using Imaris image analysis software, the filament tool was used to trace the MDN axon, as described in the *Multicolor FlpOut and quantification of morphology* section. The spots tool was then used to unbiasedly label the Brp-short staining/puncta with a 3-dimensional spot. The spots diameter was dependent on each image, but most were on average 1.3 µm. The “spots close to the filament” tool was then used to detect the number of spots located on the filament, specifically on the axon portion, with a threshold of 1. The number of spots on the axon filament was then recorded.

### Pre-synapse quantification with MDN axon varicosities

Using Imaris image analysis software, we increased the contrast within the MDN axon so the varicosities were isolated. The spots tool was then used to unbiasedly label the varicosities with a 3-dimensional spot. The average diameter was set to 1.3 µm. The surface tool was used to label the A18b neurons. The “spots close to surface” tool was then used to detect the number of spots close to the A18b surface made, with a threshold of 1. We used the Even-skipped (Eve) staining to determine the abdominal segments, as defined above.

### CaMPARI experiments

Animals were reared in the dark, with minimal light exposure during handling. At 72 hours after larval hatching, 5 animals were placed on a firm agar cap and exposed to the photoconverting light (405 nm) for 2 minutes while being monitored under a stereoscope. If the animals received a noxious head poke, they were poked in the head (mouth hooks used to determine the head) with a paint brush approximately 4-5 times, ensuring each head poke resulted in backward crawling. If the animals did not receive the noxious stimulus, they were allowed to crawl on the plate uninterrupted and observed to ensure backward locomotion did not occur during light exposure. The central nervous system was immediately dissected after photoconvertible light exposure and underwent the immunohistochemistry protocol outlined above. We stained the tissue for Flag and CaMPARI2, as previously outlined (Moeyaert et al., 2018), and found that the green signal and anti-Flag signal overlapped. We opted to only show the endogenous green (488 nm) signal in our figure panels. The CaMPARI2 signal was visualized using an AlexaFluor 555 secondary antibody. All samples were handled together and imaged on the same day to avoid batch effects.

Average intensity projections of the Red:Green signal ratio were calculated for each VNC hemi-segment. The experimental region of interest (ROI) started at the A4 A18b cell body and ended at the tip of the VNC. To calculate the background, the same ROI was shifted medially. For each signal, the background ROI fluorescence was subtracted from the experimental ROI fluorescence, before being divided by the area of the ROI. To calculate the Red:Green ratio, the background-subtracted red value was divided by the background-subtracted green value.

### Statistics

When applicable, values were normalized to the average of the control. All statistical analysis were performed with Prism 10 (GraphPad Software, San Diego, CA). Numerical data in graphs show individual measurements (dots) and means (bars). The number of replicates (n) and definition of measurement reporter (i.e. animal, axon) for each data set is in the corresponding legend.

## Acknowledgements

We thank Jim Truman for reagents, Heather Pollington and Ben Brissette for comments on this manuscript, and Heather Pollington for help making schematics in Adobe Illustrator. Antibodies obtained from the Developmental Studies Hybridoma Bank, created by the NICHD of the NIH and maintained at the University of Iowa, Department of Biology, Iowa City, IA, were used in this study. Stocks obtained from the Bloomington Drosophila Stock Center (NIH P40OD018537) were used in this study. Funding was provided by HHMI (CQD) and NICHD (F32 HD105344 to KML).

This article is subject to HHMI’s Open Access to Publications policy. HHMI lab heads have previously granted a nonexclusive CC BY 4.0 license to the public and a sublicensable license to HHMI in their research articles. Pursuant to those licenses, the author-accepted manuscript of this article can be made freely available under a CC BY 4.0 license immediately upon publication.

## Author Contributions

KML conceptualized, curated data, analyzed data, validated results, wrote, and edited the manuscript. NRC and JG curated data, analyzed data, and edited the manuscript. CQD conceptualized, supervised, wrote, and edited the manuscript.

